# An Energy Landscape Approach to Miniaturizing Enzymes using Protein Language Model Embeddings

**DOI:** 10.64898/2026.03.04.709378

**Authors:** Jakub Lála, Harsh Agrawal, Fanfei Dong, Jude Wells, Stefano Angioletti-Uberti

## Abstract

We present a general approach to find amino acid sequences corresponding to the most compact enzyme likely to retain the structure of a given catalytic site. Our approach is based on using Monte Carlo (MC) simulations to sample an energy landscape where minima correspond, by construction, to sequences with the aforementioned properties. Building on previous work (Wu et al., 2025) and with the BAGEL package (Lála et al., 2025), we implement a route to achieve this goal using only the information extracted from a protein language model (PLM), without structural information. After generating a set of candidate sequences with this PLM-guided BAGEL optimization, we further filter potential candidates for downstream experimental validation using a two-stage protocol. First, deep-learning-based structure prediction models (ESMFold, Chai-1, Boltz-2) are used to identify a structural consensus among designs with highly conserved active-site geometries, yielding many candidates with active-site RMSD below a few angstroms relative to the wild-type and pLDDT scores above 80. Second, molecular dynamics simulations are performed on a filtered subset of sequences (based on active-site RMSD and SolubleMPNN log-likelihoods) to evaluate active-site stability when including thermal fluctuations. For the most promising enzymes, these yield RMSF values in the active site below 1.0 Å and an active-site RMSD drift between 0.5 and 1.5 Å, making these mini-variants comparable to the wild type, though outcomes vary across enzymes. Given the protocol’s generality, we believe these results represent a step forward in AI-guided enzyme design. To facilitate rapid experimental validation by the broader community, we open-source all sequences generated by our computational pipeline. These include designs for four representative enzymes of this study: PETase, subtilisin Carlsberg (serine protease), Taq DNA polymerase, and VioA.

## 1 Introduction

Enzymes are ubiquitous in nature and much of biotechnology is dedicated to replicating their properties, more precisely, their ability to catalyze reactions with remarkable selectivity and high turnover [1]. Nature, guided by hundreds of millions of years of evolution, provides us with an enormous number of enzymes, many of which are either directly used in industrial settings, or serve as the starting point for optimization [2, 3]. The latter is often attempted because the natural counterpart has evolved through a set of constraints that are not necessarily similar to those encountered during their practical application in an industrial setting. For example, an enzyme might have evolved to obtain a specific product in the presence of a large mixture of potential substrates, but only requires structural stability at ambient temperature. In contrast, when used for industrial catalysis, an enzyme might encounter a more restricted set of substrates, while requiring stability at higher temperature. Similarly, the CRISPR-associated nuclease Cas9 evolved as part of bacterial adaptive immunity against phages [4], where the specificity afforded by a short guide RNA was sufficient for small bacterial genomes; in therapeutic gene editing, however, this same specificity proves inadequate for the much larger human genome, necessitating extensive engineering to reduce off-target effects [5]. These examples illustrate a broader point: natural enzymes are rarely optimal for their intended application, and re-engineering them often involves redesigning large portions of the protein.

A general desire, therefore, might be to find the shortest amino acid sequence corresponding to a specific enzymatic activity. Under the assumption that the structural geometry of a catalytic site alone determines its enzymatic activity, a natural question arises:

> *Given a specific enzyme and its known catalytic site geometry, what is the shortest amino acid sequence that will fold in such a way as to preserve the geometry of the active site, and thus preserve its catalytic activity?*

In protein design broadly, this question is known as the *motif scaffolding* problem. An answer to this question is not purely of academic interest, but instead has various practical implications [6]. For example, an enzyme’s size limits its possibility to be delivered via viral vectors or nanoparticles, preventing its use for nanomedicine applications [7]. Similarly, larger enzymes are more difficult to manufacture at scale in high yield, again limiting their use [6, 8]. Although the evolutionary pressure to reduce genome size might have driven biology to produce very compact enzymes, many of these may remain multifunctional and thus such pressure might not have found the shortest possible sequence. Deep-learning models for proteins now offer the tools to systematically search for such optimally compact sequences, motivating the present study.

In this manuscript, we build on the previous work by Wu et al. [9] and Lála et al. [10], and introduce a general approach to miniaturizing enzymes. We do so by carrying out a Monte Carlo (MC) sampling of an energy landscape, specifically a grand-canonical potential energy determined by two sequence-dependent terms: a) an embedding similarity energy *E*_embedding_ guiding the search to conserve the semantic embedding of the active-site residues and their surrounding buffer (which together we refer to as the *immutable* region) as close as possible to their embedding within the full-length wild type (WT) sequence (as in Wu et al. [9]), and b) a chemical potential energy *E*_chemical_ penalizing for longer sequences. Throughout this paper, we use “active site” as shorthand for the full immutable region for simplicity, although the immutable region strictly includes both the crucial catalytic residues and a surrounding sequence buffer (see Methods 4.1 for details).

This is formally defined as:

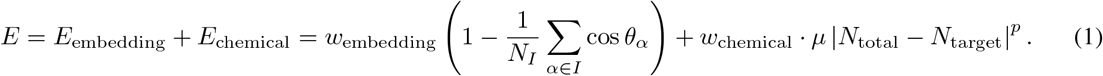

*E*_embedding_ is defined over the immutable residue group *I*, containing *N*_*I*_ = | *I* | residues. For each residue *α*, we compute the angle *θ*_*α*_ between two high-dimensional embedding vectors of the current and reference sequences, *e*_*α*_ and 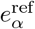 respectively, such that 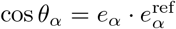 corresponds to their cosine similarity. These can be obtained from an embedder of choice, in our case, a PLM. The chemical potential term *E*_chemical_ penalizes deviations from a target sequence length, where *µ* is the chemical potential, *N*_total_ is the total number of residues in the system, *N*_target_ is the target number of residues, and *p* is an exponent controlling the sharpness of the length penalty. *w*_embedding_ and *w*_chemical_ are user-defined weights to be balanced out for the respective terms based on specific application.

Casting the problem in statistical mechanics terms, the product *w*_chemical_ · *µ* = *µ*\* can be identified as the effective chemical potential of a grand-canonical ensemble [11], with ⟨| *N*_total_ − *N*_target_ |^*p*^⟩ as its conjugate variable at inverse temperature *β* = 1*/*(*k*_*B*_*T*). This analogy provides a principled mechanism for controlling sequence length: MC sampling at temperature *T* minimizes the grand-canonical potential, and increasing *µ* systematically drives the algorithm toward shorter candidate sequences.

In a more practical sense, we implement the algorithm using the `BAGEL` software [10]. First, we test various values for the strength of the chemical potential energy *w*_chemical_, and validate (albeit *in silico*) the designed mini-variants with deep-learning folding algorithms and, furthermore, with molecular dynamics (MD) to include thermal fluctuations in our analysis and, more generally, to employ an orthogonal, physics-based method. We focus on four different enzymes: PETase (PDB ID: 5XJH, referred to as *petase*), subtilisin Carlsberg (PDB ID: 1SBC, referred to as *protease*), Taq DNA polymerase (PDB ID: 1TAQ, referred to as *taq*), and VioA (PDB ID: 6FW9, referred to as *vioA*), and provide the final sequences from our computational pipeline in an open-source Zenodo repository. Details on the enzymes are provided in Methods 4.1.

## 2 Results

### 2.1 Strength of miniaturization can be controlled by chemical potential energy

As an initial validation of the approach, and to justify the choice of the initial parameters for a given temperature at which we plan to sample new mini-variants, we carry out an initial sweep over different weights *w*_chemical_ for the chemical potential energy *E*_chemical_. This determines the strength of miniaturization, i.e., how much we penalize for longer sequences. A few expected general patterns appear as we increase the weight *w*_chemical_, summarized in Figure 2. Even from a visual inspection of Figure 2A, it is clear that shorter sequences come at the cost of loss of active-site structure: larger values of *w*_chemical_ drive greater size reduction but increasingly disrupt the immutable region, likely resulting in a complete loss of enzymatic activity. The remaining enzymes are visualized in Supplementary Information (Figure 7). This is reflected quantitatively in lower mean predicted Local Distance Difference Test (pLDDT) scores and higher Root Mean Squared Deviation (RMSD) values computed on the immutable region *I* alone in Figure 2B. At lower values of *w*_chemical_, this effect becomes less pronounced: *petase, protease*, and *taq* retain high pLDDT and low RMSD values across *w*_chemical_ ∈ {0.001, 0.0005, 0.0001}, while *vioA* shows a more gradual drift. Nevertheless, regardless of the weight *w*_chemical_ employed, we observe both a reduction in sequence length and a diversification of sequences away from the WT (Table 1). For details of the method, see Methods 4.2.1. Given these results and for simplicity, we choose *w*_chemical_ = 0.001 for all enzymes. We note, however, that the optimal choice is system-dependent: *vioA* exhibits a gradual degradation across weights, whereas *protease* displays a sharp transition between *w*_chemical_ = 0.001 and 0.0005, with a pronounced gap in the pLDDT distributions suggesting a threshold-like regime.

**Figure 1:**
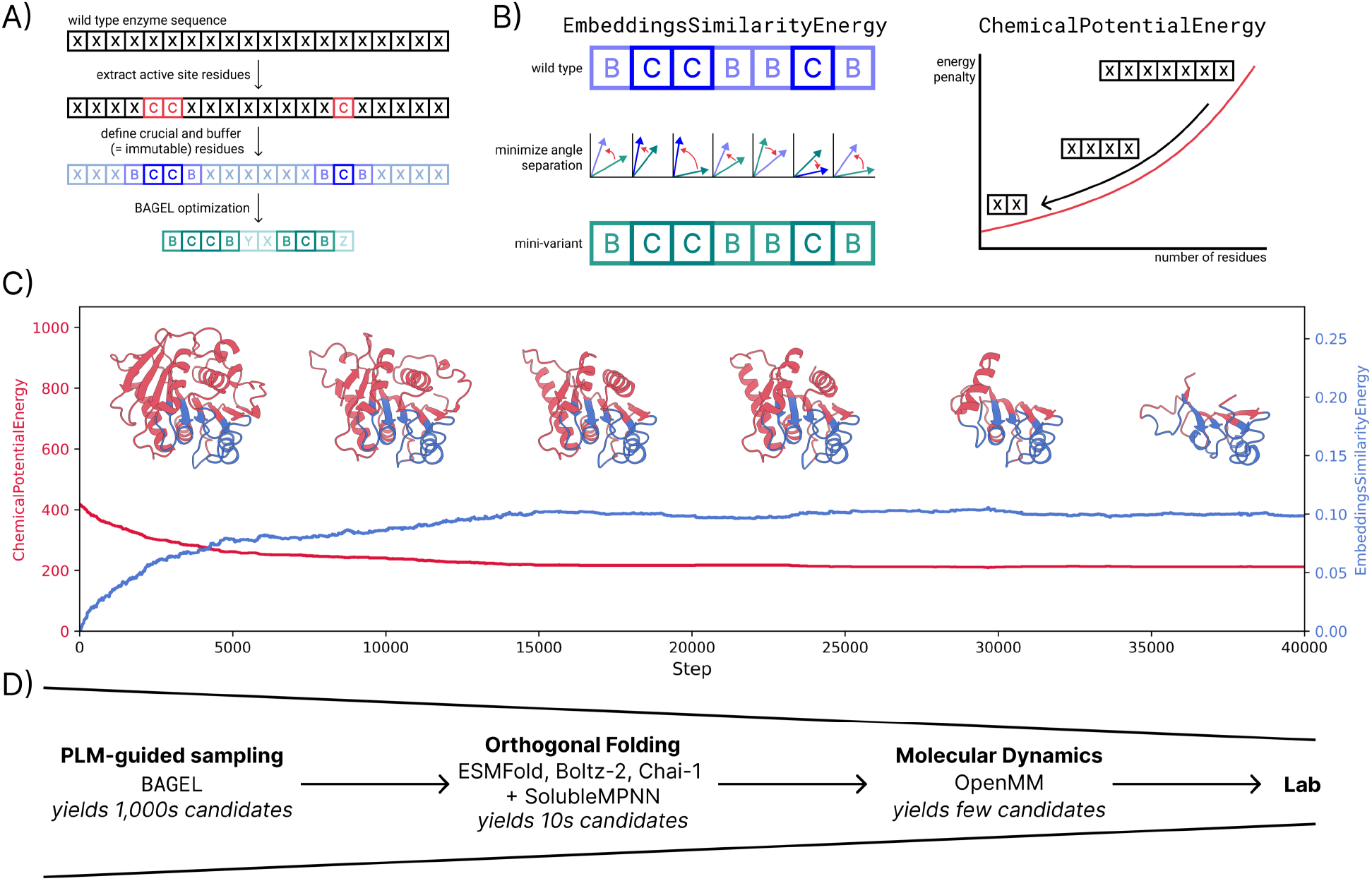
Overview of the PLM-guided enzyme miniaturization pipeline. A) Graphical summary of our approach. After extracting *crucial* active-site residues (C) from the WT sequence, we identify a surrounding buffer sequence *±* a few residues (B) on either side of the crucial ones. The whole part is defined as an *immutable* region (*I* = B ∪ C), which remains fixed during the optimization. Due to the form of the energy, see Equation 1, the optimization (i.e. energy minimization) leads to `BAGEL`-guided [10] sequence miniaturization. B) `BAGEL` energy terms. We employ previously defined `EmbeddingsSimilarityEnergy` (*E*_embedding_) [9], where the embeddings in the immutable region are compared between the mini-variant and the WT. Additionally, we introduce a new `ChemicalPotentialEnergy` (*E*_chemical_) to drive the optimization towards shorter sequences by penalizing sequence length, and thus achieving miniaturization. C) `BAGEL` evolution of a system. After an initialization at the WT reference, the MC optimization iteratively samples substitutions, deletions, and insertions of amino acids. *E*_embedding_ starts at zero, i.e., at the WT reference, and then gradually reaches a plateau after a transient region. *E*_chemical_ monotonically decreases until a similar plateau is reached and the sequence-length penalty is no longer strong enough to shrink the enzyme further. These plateaus occur when the two energy terms are balanced, i.e., reducing the sequence length through *E*_chemical_ is no longer beneficial enough to offset the loss of similarity in *E*_embedding_. Note, these energy terms are dimensionless. D) Full funnel workflow. PLM-guided MC sampling generates thousands of candidate sequences, which are progressively filtered using folding algorithms and molecular dynamics simulations to yield a tractable set for downstream experimental validation.

**Figure 2:**
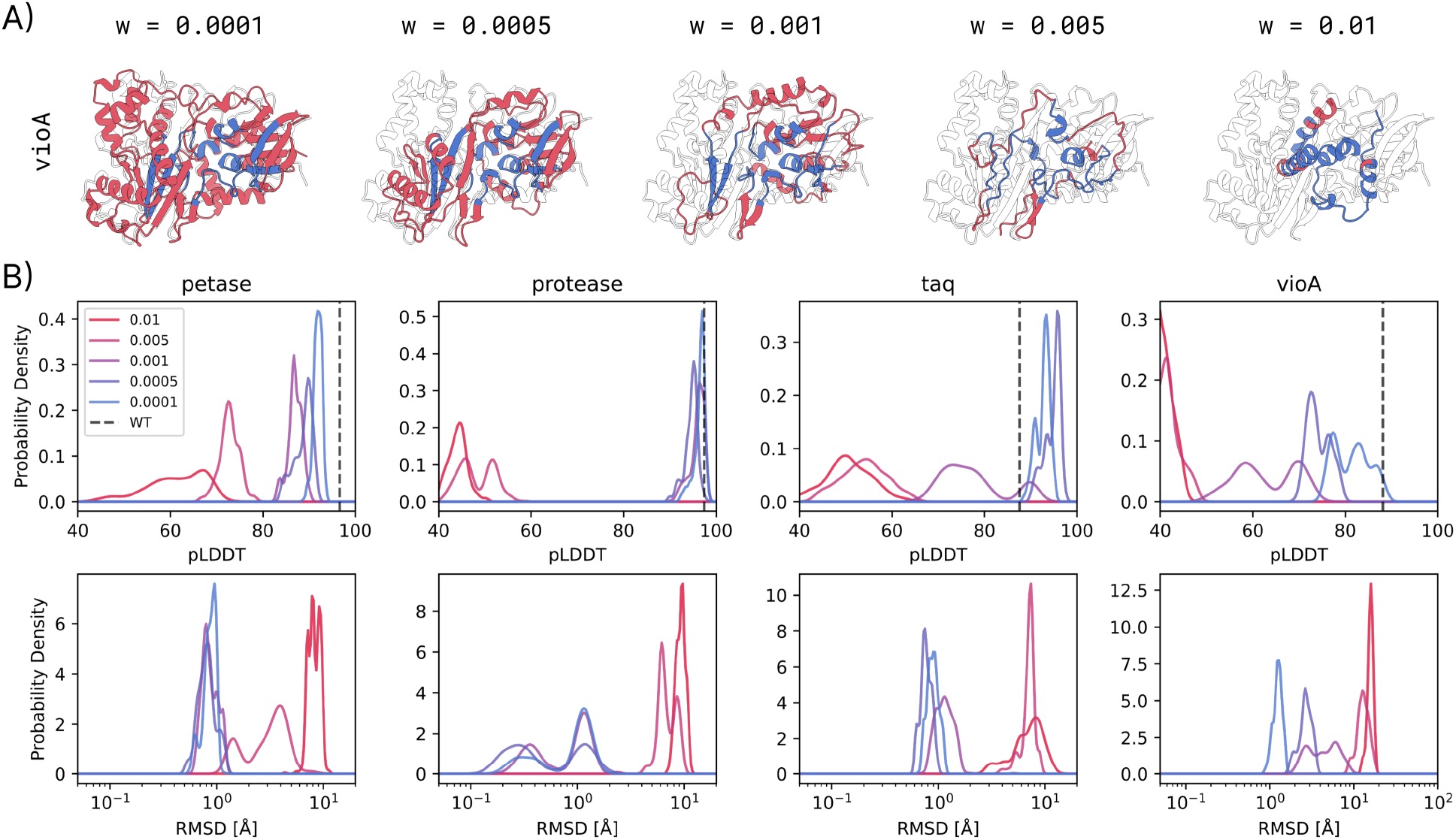
Effect of the chemical potential weight on PLM-guided enzyme miniaturization. A) Representative *vioA* mini-variant structures from `BAGEL` [10] optimizations at increasing weights *w*_chemical_, folded with ESMFold [12]. The immutable region (*I*) is colored in blue, the remaining sequence in red, and the WT structure is shown as a black silhouette. As *w*_chemical_ increases, sequences become progressively shorter and the immutable region becomes structurally compromised. B) Distributions of pLDDT (top) and immutable-region heavy-atom RMSD (bottom) across sampled mini-variant sequences for each enzyme, colored by *w*_chemical_ (see legend). The dashed vertical line indicates the WT pLDDT. RMSD is computed against the WT structure folded with ESMFold. RMSD densities are computed in log_10_-space.

**Table 1:** Enzyme miniaturization outcomes across varying chemical potential weights. Results of sampled mini-variants across different weights *w*_chemical_. By increasing *w*_chemical_, the final sequence length is smaller, while having a large Levenshtein distance [13], i.e., sequence dissimilarity, to the full-length WT sequence.

| Enzyme | WT | $w_{\text{chemical}}$ | | | | |
| --- | --- | --- | --- | --- | --- | --- |
|  |  | 0.0001 | 0.0005 | 0.001 | 0.005 | 0.01 |
| petase | **263** | 0.84 | 0.69 | 0.68 | 0.45 | 0.31 |
| protease | **275** | 0.83 | 0.76 | 0.74 | 0.43 | 0.27 |
| taq | **823** | 0.31 | 0.27 | 0.22 | 0.11 | 0.09 |
| vioA | **417** | 0.98 | 0.65 | 0.50 | 0.32 | 0.24 |

| Enzyme | $w_{\text{chemical}}$ | | | | |
| --- | --- | --- | --- | --- | --- |
|  | 0.0001 | 0.0005 | 0.001 | 0.005 | 0.01 |
| petase | 0.62 | 0.68 | 0.67 | 0.81 | 0.89 |
| protease | 0.47 | 0.53 | 0.55 | 0.76 | 0.86 |
| taq | 0.80 | 0.82 | 0.85 | 0.92 | 0.94 |
| vioA | 0.71 | 0.75 | 0.78 | 0.86 | 0.92 |

### 2.2 Embedding-based approach proves effective at certain sequence lengths

Before proceeding, we notice that for *petase* and *protease*, the mini-variants end up being trimmed primarily at the N- and C-termini up until the immutable region. This makes us speculate that the most naive (but potentially also effective) approach to reducing the size of an enzyme is by deleting the residues at either ends in a pre-defined algorithmic way, without necessarily needing to compensate with substitutions elsewhere in the sequence to stabilize the miniaturized fold. Therefore, we establish such a baseline experiment on the four enzymes in Figure 3 in a two-stage strategy (detailed in Methods 4.3, Algorithm 1): initially, we alternately remove residues from the N- and C-termini until one end reaches the immutable region, followed by round-robin deletions of all remaining mutable segments.

**Figure 3:**
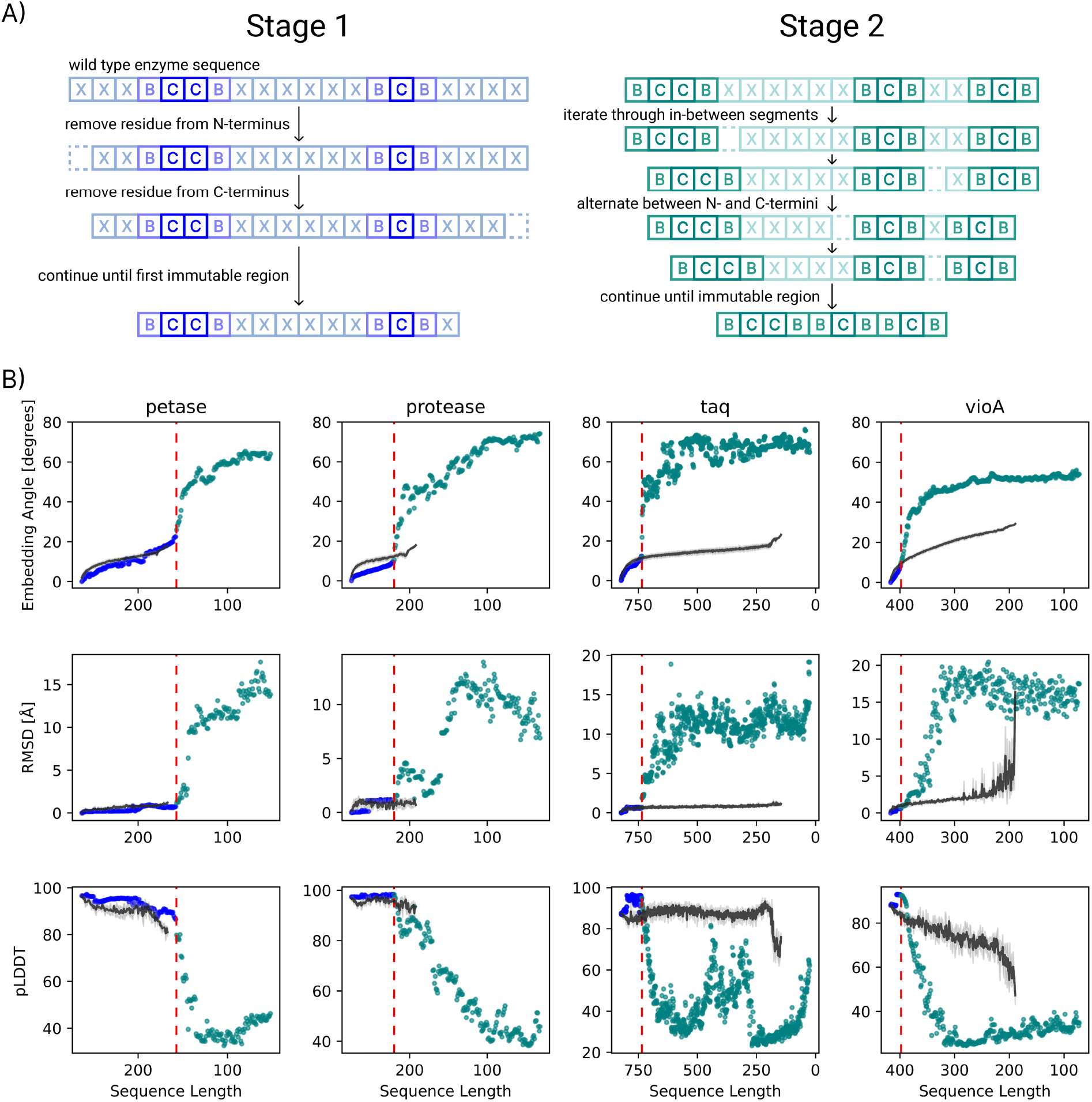
Comparison of naive trimming with PLM-guided BAGEL optimization for enzyme miniaturization. A) Schematic of the two-stage manual trimming baseline. Mutable residues (X) are progressively removed while immutable residues (B = buffer, C = crucial) are preserved. In Stage 1, residues are alternately removed from the N- and C-termini until either end reaches the immutable region. In Stage 2, all remaining mutable segments — including any leftover terminal residues and segments between distinct immutable regions — are trimmed in round-robin order, alternating from each end within each segment, until only immutable residues remain. B) Comparison of the manual trimming trajectory to `BAGEL`-sampled [10] sequences for the four enzymes. Rows show, from top to bottom: embedding angle (degrees), RMSD (Å), and pLDDT. Blue marks Stage 1, teal marks Stage 2, the red dashed line indicates the transition between them, and black marks `BAGEL`-sampled [10] sequences at corresponding sequence lengths. The x-axis runs from WT length (left) to zero (right).

In Figure 3B, we compare this manual trimming trajectory to the trajectory produced by the `BAGEL` optimization (from a production run, detailed in Methods 4.2.2). In general, we observe a two-stage behavior: at first, the `BAGEL`-derived sequences are comparable to the manually miniaturized enzymes, as we only remove residues at either termini (both in terms of embeddings, and structural metrics); but once we need to remove internal residues, `BAGEL` greatly outperforms the manual trajectory. This is less pronounced for *petase* and *protease*, where `BAGEL` does not find shorter sequences beyond the length defined by the first stage. On the other hand, for *taq* and *vioA*, we observe that at shorter sequences, there is a much smaller embedding angle (and thus also *E*_embedding_), much lower RMSD, and much higher pLDDT in the immutable region, compared to the manual trajectory. This highlights `BAGEL`’s ability to find substitutions, such as to i) keep the embeddings of the immutable region aligned to that of the WT, ii) conserve the geometry of the region in terms of RMSD, and iii) retain the confidence of the predicted structure, despite not modeling the structure explicitly in the PLM. Thus, we see a confirmation that when we achieve a smaller embedding angle than the manual trajectory (top row in Figure 3B), we can assume greater structural agreement of the mini-variant to the WT (middle row in Figure 3B).

### 2.3 BAGEL finds miniaturized variants with active-site geometries akin to the wild type

To avoid adversarial examples, i.e., sequences a model folds confidently but incorrectly, we fold all mini-variants (produced from the production run, see Methods 4.2.2) with three independent structure prediction models, namely ESMFold [12], Chai-1 [14] and Boltz-2 [15] (see Methods 4.4). The choice of these models was dictated by both practical considerations (the use of open-source models without licensing restrictions) and the desire to maximize the diversity in model architectures (e.g., in how initial per-residue embeddings are generated, or what structure module is employed). Therefore, by achieving a consensus among these models, we aim to minimize the number of potentially adversarial sequences.

Figure 4 summarizes the results. In general, *petase* and *protease* show close agreement at low RMSD values and high pLDDT scores across all models. For *taq*, this agreement is worse, with Boltz-2 yielding higher RMSD, while having comparable pLDDT to ESMFold. *vioA* then shows both the largest deviation from the WT, and the lowest consensus among the three models, both in terms of RMSD and pLDDT. We attribute this to an overly large *w*_chemical_ for this particular enzyme. Furthermore, we observe an inverse relationship between RMSD and pLDDT, i.e., designs closer to the WT tend to be predicted with higher confidence; this potentially obvious pattern is not a general necessity, but is supported by the analysis in Supplementary Information C (Figure 8).

**Figure 4:**
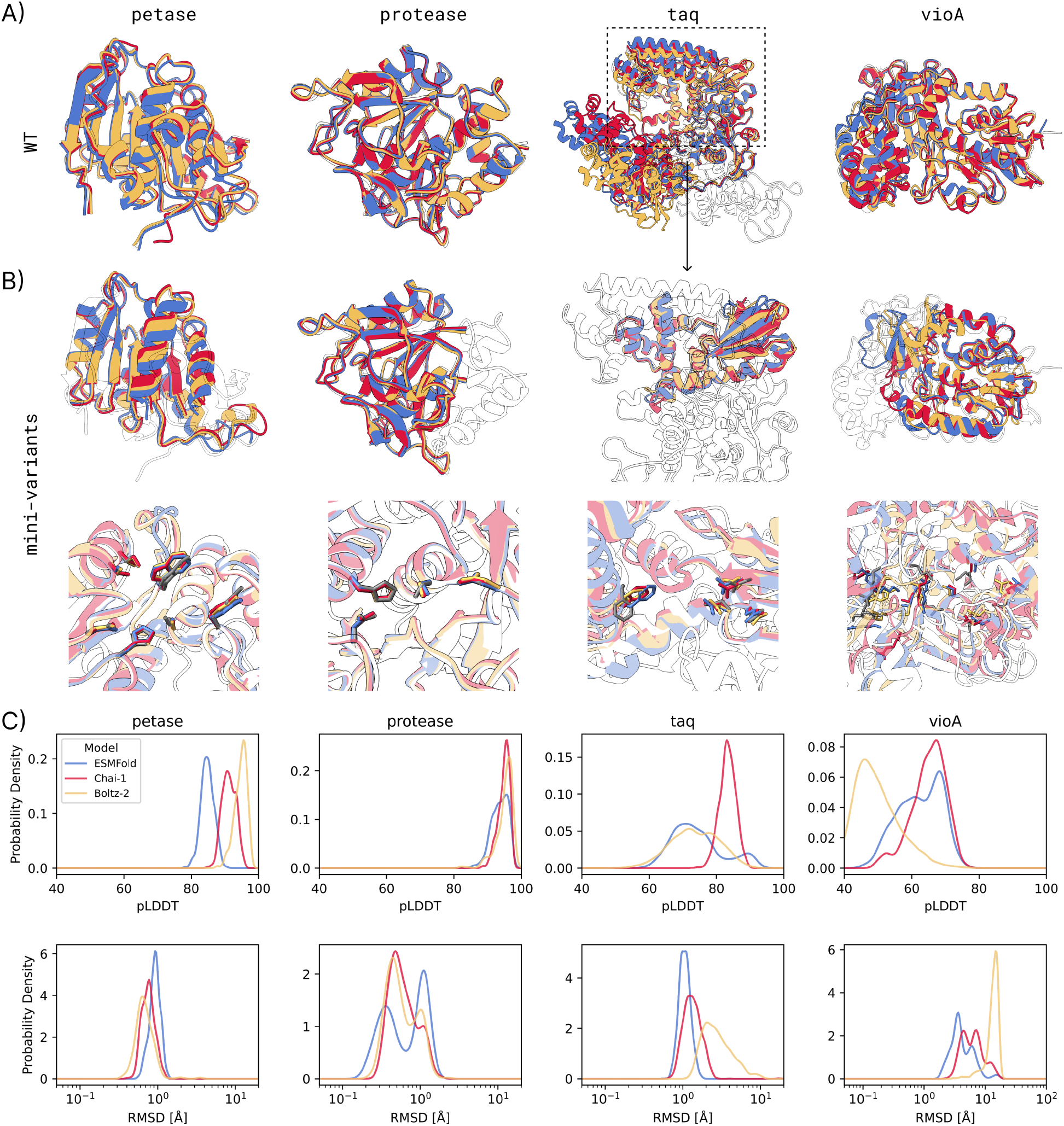
Multi-model structural validation of miniaturized enzyme variants. A) Visualization of the WT sequences folded with different models (ESMFold [12], Chai-1 [14], Boltz-2 [15]), overlaid with the crystal structure (black outline). We see an agreement for all enzymes apart from a region in *taq*, where the model predictions differ from the crystal structure by a relative positioning of the two domains. B) Visualization of the best mini-variant sequences per enzyme folded with each model, overlaid with the WT crystal structure (black outline). The best mini-variant is the one with the lowest average RMSD (of the immutable region) across all models when compared to the folded WT sequence (with each model). The upper row shows the full structure, the lower row shows a close-up on the crucial active-site residues (*C*). *petase* and *protease* show close agreement with the WT, while *taq* shows more deviation (with a wrong rotation on the left residue), and *vioA* with its large active site shows even more deviation across most residues. C) Distribution of pLDDT and immutable region RMSD (heavy atoms) for the mini-variants. Probability density distributions are shown for sequences folded with ESMFold (blue), Chai-1 (red), and Boltz-2 (yellow). RMSD densities are computed in log_10_-space. Note that according to all models, there is a large fraction of sequences (apart from *vioA*) folding with very high confidence and low RMSD for the catalytically important region.

Note we compute RMSD by comparing each mini-variant to the WT folded by the same model, rather than to the crystal structure. This avoids penalizing mini-variants for discrepancies that arise from a model’s imperfect reconstruction of the WT itself. Our goal is thus to find mini-variants that align with the folding models’ internal representation of the functional enzyme, rather than necessarily reproducing the crystal structure; experimental validation of this hypothesis is left for future work. After filtering by RMSD consensus across the three models, and also ranking by SolubleMPNN [16] log-likelihood (Methods 4.4), we select the top-16 candidates per enzyme for MD validation.

### 2.4 Filtered mini-variants show some dynamical agreement to the wild type

Folding algorithms do not model protein dynamics with physics-based forces, nor are they able to sample different, potentially competing structures according to the physically meaningful Boltzmann distribution that the system is supposed to follow. For this reason, to quantify the effect of thermal fluctuations, we perform MD simulations (details in Methods 4.5). Besides gauging these effects, MD also provides an orthogonal, independent way to evaluate the stability of the active-site geometry.

We first examine the per-residue thermal fluctuations by computing the Root Mean Square Fluctuations (RMSF):

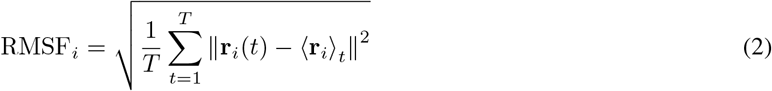

where **r**_*i*_(*t*) is the position of the Cα atom of residue *i* at frame *t, T* is the total number of frames in the trajectory, ⟨ **r**_*i*_ ⟩_*t*_ is the time-averaged position of that atom computed over the trajectory, and ∥ · ∥ denotes the Euclidean norm. For each entry – whether the WT or a mini-variant – every frame is first superimposed onto the first frame of its own trajectory in terms of all of the *C*_*α*_ atoms. Figure 5 shows the best-performing mini-variant based on this RMSF metric, with Figure 10 in the Supplementary Information providing the data for all the 16 mini-variants simulated with MD. Overall, the mini-variants do not replicate the WT profile exactly, but they remain stable and maintain comparable local flexibility at the active site despite being generated solely from PLM guidance. Outside of the immutable regions, the RMSF might be large, but we neither filter based on that, nor can we properly compute it given that the mapping between residues in the mini-variant and the WT is non-trivial as the global folds differ. For instance, we see that both *petase* and *protease* have a newly exaggerated peak at residues 120, and 100 or 160 respectively, where a former hinge in the WT, i.e., an increased thermal fluctuation, is much more pronounced in our mini-variants. To complement this global measure with an active-site deviation over the trajectory, we compute an RMSD-like quantity of the heavy atoms in the immutable residue group *I* at each frame *t*:

**Figure 5:**
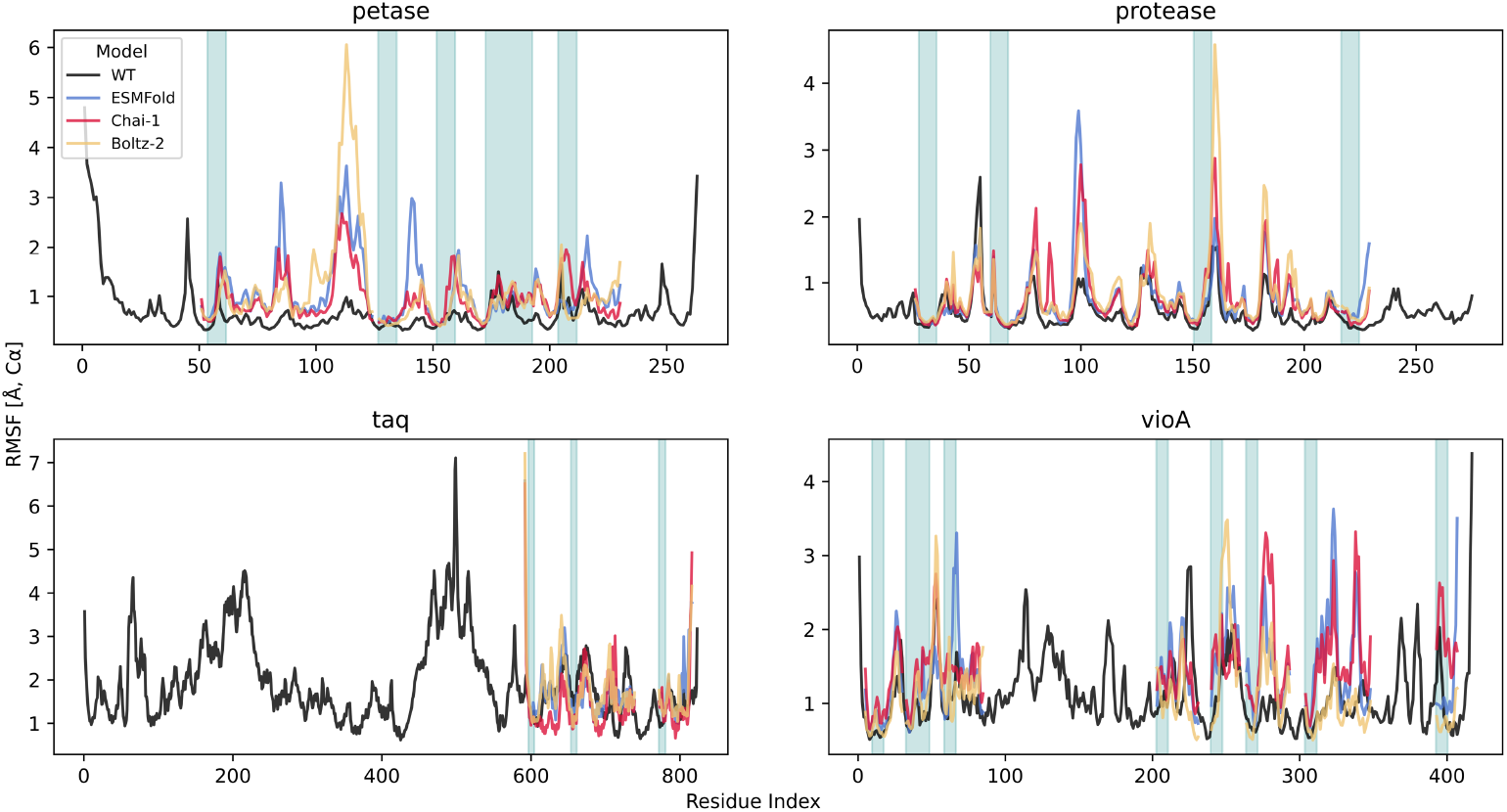
Molecular dynamics validation of the best miniaturized enzyme variants. Per-residue Cα RMSF (defined in Equation 2) profile of the best-performing mini-variant for each enzyme, compared to the full-length WT. The best mini-variant per enzyme was selected by the smallest average deviation from the WT RMSF profile, yielding mini-petase-138, mini-protease-35, mini-taq-1547, and mini-vioA-980. WT is shown in black, while mini-variant simulations are shown for initializations from ESMFold [12] (blue), Chai-1 [14] (red), and Boltz-2 [15] (yellow). Shaded teal regions indicate the immutable residues.

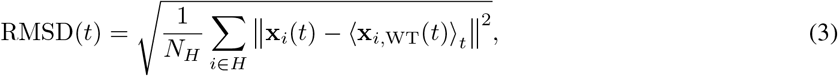

where *H* is the set of heavy atoms across all residues in *I* (defined in Equation 1), *N*_*H*_ = | *H* |, **x**_*i*_(*t*) is the position of atom *i* ∈ *H* at frame *t* (after superimposing the mutant immutable region onto the WT immutable region’s average structure), and ⟨ **x**_*i*,WT_(*t*) ⟩_*t*_ is the time-averaged position of that atom in the WT trajectory. Figure 6 shows that some of the mini-variants are in close agreement in terms of their RMSD deviation from the average active-site geometry in the WT, while others deviate significantly. This means that in the best cases, the immutable region deviates only by 1–2 Å at most, on par with the deviation for the WT itself.

**Figure 6:**
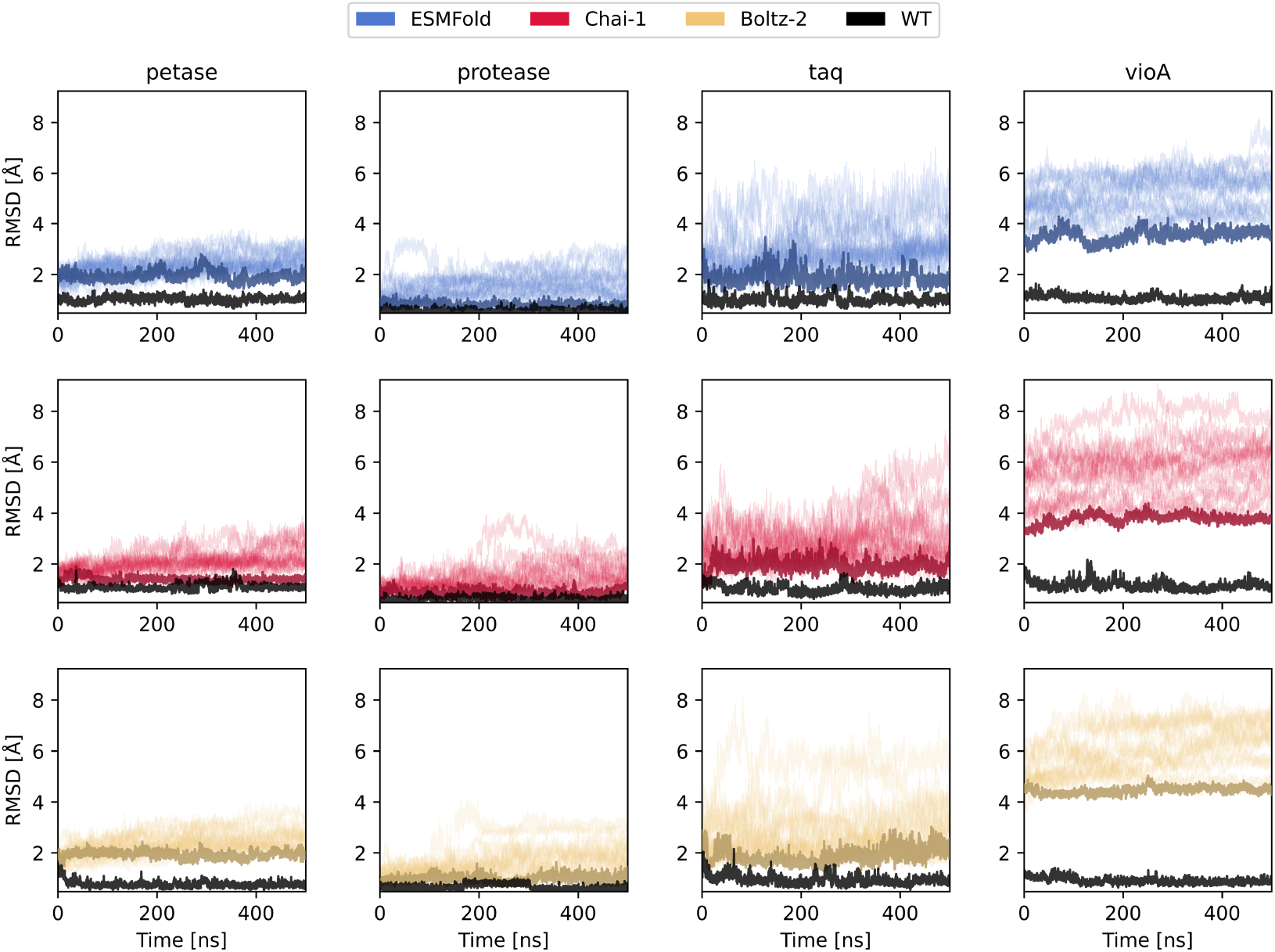
Active-site stability of miniaturized enzyme variants over MD trajectories. Immutable region heavy-atom RMSD (defined in Equation 3) of mini-variants compared to WT over time. The RMSD trajectory is computed against the average structure of the WT trajectory. Rows correspond to starting structures folded by ESMFold [12] (blue), Chai-1 [14] (red), and Boltz-2 [15] (yellow), while columns correspond to the four enzymes. Each panel shows the WT (black) alongside the mini-variants for that folding model. WT is initialized from folded structures, not the crystal one. The best mini-variant, defined as the one with the lowest mean RMSD over the trajectory, is highlighted in a darker shade.

## 3 Discussion

By implementing a chemical potential energy term together with a grand-canonical sampling minimizer in `BAGEL` [10], we have shown that a PLM-guided approach can miniaturize enzymes while constraining the embeddings of the active site residues and, as a result, conserving the local geometry of the catalytic site. In practice, this amounts to moving from a canonical to a grand-canonical ensemble, exploiting the well-known relation between the average system size and the chemical potential. Under the assumption that such per-residue PLM embeddings capture local sequence-context features, we speculate that the catalytic activity should also be retained. This hypothesis is grounded in the observation that, during evolution, proteins with similar functionality typically conserve the overall fold and local geometry of the active region [17, 18]. Following the principle that structure implies function, conservation of the catalytic-site geometry should bias sequence generation towards variants that retain catalytic activity — though experimental validation will be needed to confirm this, and outcomes will likely differ across enzymes. Indeed, one might argue that agreement in the active-site fluctuations, rather than in the static geometry alone, provides an additional layer of support for retaining catalytic function.

The four enzymes studied here exhibit markedly different outcomes, reflecting both the method’s capabilities and its limitations. *petase* and *protease* show the most promising results, with mean final sequence lengths of 177 and 203 amino acids respectively (33% and 26% reduction from WT), low RMSD values (below 1.0 Å), and high confidence scores across all three folding models (namely, ESMFold, Chai-1, and Boltz-2). However, we note that the miniaturization in these cases was achieved primarily through terminal trimming, with limited removal of internal residues between immutable regions, though substitutions in those regions are still carried out by the optimization. Our manual trimming analysis demonstrated that `BAGEL`’s advantage becomes most apparent when internal deletions are necessary, suggesting that *petase* and *protease* may represent relatively straightforward miniaturization targets.

In contrast, *taq* and *vioA* demonstrate more dramatic miniaturization, with mean final lengths of 172 and 205 amino acids (79% and 51% reduction, respectively). For *taq*, the massive reduction involved removal of the N-terminal domain containing the 5’ nuclease activity, thereby retaining only the C-terminal polymerase domain where the catalytic residues (Asp610, Phe667, Asp785, Glu786) reside. We note that the WT Taq polymerase’s 3’-5’ exonuclease domain is already non-functional [19], and removal of the 5’ nuclease domain may be acceptable for applications requiring only DNA polymerase activity, though it would preclude use in probe-based qPCR assays that rely on the 5’ nuclease function [20]. Despite this dramatic change, the mini-taq variants maintain reasonable *in silico* metrics, with RMSD values below 1.5 Å for candidates folded with ESMFold and Chai-1, though Boltz-2 yields higher RMSD for this enzyme.

The *vioA* results present the greatest uncertainty. VioA is a two-domain FAD-dependent oxidase, and the 51% size reduction resulted in structures with substantially higher RMSD values (requiring a 5.5 Å filtering threshold compared to 1.0 Å for *petase* and *protease*) and lower confidence scores across all folding models. We note that *mini-vioA* shows large structural deviations in MD, raising questions about whether these variants would retain the complex FAD-binding pocket and oxidase activity. These differential outcomes highlight that miniaturization feasibility is enzyme-dependent, and thus we invite the community to employ this `BAGEL` recipe on their enzyme of interest.

Our approach differs fundamentally from structure-based motif scaffolding methods such as RFDiffusion [21–23], which generate new scaffolds around a predefined geometric motif. It instead works directly in sequence space, utilizing the ever-improving power and generalizability of PLMs without requiring 3D structural input. Furthermore, RFDiffusion does not natively support miniaturization, so a user would need to sweep over scaffold lengths to identify the smallest viable design – a search that `BAGEL` handles naturally through the strength of the chemical potential energy term.

More directly related to our goal is Raygun [24], which specifically targets protein miniaturization and has demonstrated impressive results with experimental validation for green fluorescent protein. Nevertheless, their approach has several limitations. First, the user cannot specify which residues to explicitly conserve. With a growing body of evidence of which residues play a critical role in catalysis, this would require new models, while `BAGEL` has this feature directly built-in, allowing the user to make any specific residues immutable. Secondly, the authors suggest fine-tuning Raygun’s decoder to avoid catastrophic forgetting and match the decoded sequences to the enzyme group, while our approach does not require any additional data at all. This could in principle allow the design of *de novo* enzymes, or proteins with very low number of homologs. Thirdly, Raygun requires specification of a target length *a priori*, something difficult to gauge from a rational perspective. Although `BAGEL` also requires a user-defined parameter for the strength of miniaturization, that being the weight of the chemical potential energy term, a few reasonable values across several orders of magnitude can be swept, as shown in Section 2.1, with the optimization left to evolve and balance out the amount of *squeezing* allowed before the semantic context of the active site is lost in the embeddings.

In our protocol, we first validated all candidates with orthogonal deep-learning folding algorithms, reserving the more computationally expensive molecular dynamics simulations for a filtered subset. This was to provide further evidence before experimental validation, as one can test only a handful of sequences, compared to thousands or more produced by the MC sampling process. We note, however, that this filtration remains to be one of the key open problems in current protein design pipelines. Therefore, we have deliberately avoided pursuing extensive computational scoring and ranking beyond our multi-model structural consensus and MD stability assessment, as there remains insufficient evidence in the literature that scoring functions or even physics-based force fields can accurately predict experimental success rates. For instance, our mini-variants exhibit lower SolubleMPNN log-likelihood scores than their wild-type counterparts (Supplementary Information D), which we attribute to sequence novelty rather than definitive insolubility: these *de novo*-like sequences likely lie outside the training distribution of models fitted to natural proteins, and thus low likelihood may reflect unfamiliarity rather than poor biophysical properties. We thus await experimental validation before further analyzing and correlating such *in silico* scores to experimental hit rate.

Importantly, the generality of our approach makes it applicable to diverse enzyme classes without requiring system-specific training. The four enzymes studied here represent distinct catalytic mechanisms and structural folds, suggesting broad applicability. Unlike approaches that require fine-tuning on enzyme-specific datasets, our PLM-based framework can be directly applied to understudied or *de novo* enzymes where homologous training data is scarce, making it particularly valuable for emerging biotechnology applications.

Looking forward, several promising extensions could further improve this approach. Hybrid optimization procedures that integrate PLM sampling with expensive folding oracle and MD simulation steps could provide stronger guarantees of structural integrity. Emulators of protein fluctuations, such as BackFlip [25], could replace prohibitively expensive MD simulations while retaining physics-informed validation. Indeed, in our mini-variant systems, we do observe the previously described correlation between pLDDT and RMSF [26] (Figure 11 in Supplementary Information), suggesting that one could potentially rely on folding models alone, reserving MD simulations only for a filtration step, instead of directly incorporating them into the energy function. Additionally, one could initialize the optimization with the immutable region alone instead of the WT sequence, and thus allow the sequence to *grow* instead of being *squeezed*. Apart from being interesting from an academic sense, this could offer inference speed-up benefits when structure is modeled explicitly, as shorter sequences fold faster. More broadly, because `BAGEL` delegates sequence evaluation to external models, it will naturally benefit from improvements in PLM and structure prediction models without requiring any algorithmic changes.

Lastly, we do not claim that these mini-variants will yield comparable (or, for that matter, higher) catalytic rates than their full-sequence parents. Instead, we showcase a promising inference-time method for employing the already powerful PLMs to a long-standing problem in biotechnology. To facilitate and speed up experimental verification by the whole community, we are open-sourcing all of the data from our pipeline, detailed in the Code and Data Availability section. Thanks to the large-scale pre-training of open-sourced models, we hope this method sparks new ideas of how `BAGEL`’s modules can be repurposed for diverse, user-defined applications.

## 4 Methods

### 4.1 Enzyme Extraction

We focus on four enzymes (PDB IDs listed in Table 2): PETase, subtilisin Carlsberg, Taq DNA polymerase I, and the FAD-dependent L-tryptophan oxidase VioA. PETase is a PET-depolymerizing hydrolase central to enzymatic plastic recycling, making it a high-impact target where miniaturization could yield smaller, more engineerable depolymerases for scalable PET bioconversion [27]. Subtilisin Carlsberg is a canonical industrial serine protease (notable in detergent formulations), so a miniaturized variant would provide a compact, robust proteolysis module with immediate bioprocess relevance [28]. Taq DNA polymerase is the workhorse thermostable polymerase underpinning PCR-based amplification used throughout molecular biology and diagnostics, motivating miniaturized variants for more portable and engineerable nucleic-acid processing modules [29]. VioA initiates violacein biosynthesis via FAD-dependent L-tryptophan oxidation, providing a tractable testbed for miniaturizing redox enzymes relevant to metabolic engineering of bioactive natural products [30].

**Table 2:**
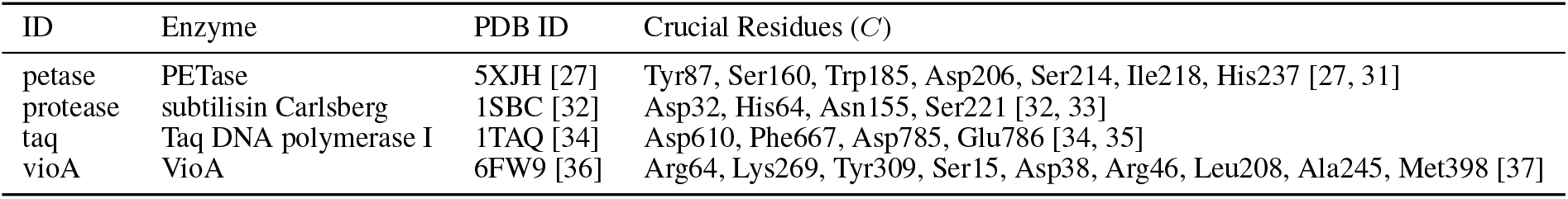
Crucial residues defining the immutable region for each enzyme. Crucial residues (*C*) for each enzyme, retrieved from literature. The full immutable region *I* = *B* ∪ *C* additionally includes a buffer of 4 residues to the left and 3 to the right of each crucial residue. For VioA, FAD-binding residues were identified by visual inspection of the crystal structure.

| ID | Enzyme | PDB ID | Crucial Residues ($C$) |
| --- | --- | --- | --- |
| petase | PETase | 5XJH [27] | Tyr87, Ser160, Trp185, Asp206, Ser214, Ile218, His237 [27, 31] |
| protease | subtilisin Carlsberg | 1SBC [32] | Asp32, His64, Asn155, Ser221 [32, 33] |
| taq | Taq DNA polymerase I | 1TAQ [34] | Asp610, Phe667, Asp785, Glu786 [34, 35] |
| vioA | VioA | 6FW9 [36] | Arg64, Lys269, Tyr309, Ser15, Asp38, Arg46, Leu208, Ala245, Met398 [37] |

For each of these, we identify from the literature the *crucial* residues (*C*) known to play a direct role in enzymatic activity. Nevertheless, to ensure we retain activity and do not accidentally remove surrounding residues that might be essential (but, for instance, have not been proven in the literature yet to be critical), we also add an effective “buffer” zone (*B*) of 4 residues to the left and 3 residues to the right of any crucial residue in the sequence. Together, the crucial residues and their buffer form the *immutable* region (*I* = *B* ∪ *C*, as in Equation 1) that we do not mutate during the optimization. All RMSD and pLDDT metrics reported throughout the paper are computed over this full immutable region *I*, not only the crucial residues *C*. Table 2 shows the exact residue positions and identities of the crucial residues considered.

### 4.2 Monte Carlo Optimization in BAGEL

Similar to Wu et al. [9], we employ an `EmbeddingsSimilarityEnergy`, *E*_embedding_, from `BAGEL` [10], but also add a penalty for growing sequence length with `ChemicalPotentialEnergy`, *E*_chemical_. Thus, we can let the sequence length change arbitrarily as the energy landscape is traversed, by adding insertions and deletions using the `GrandCanonical` mutation protocol. All embeddings are computed using ESM-2 (`esm2_t33_650M_UR50D`) [12]. The embedding similarity weight is fixed at *w*_embedding_ = 1.0 throughout all experiments, while *w*_chemical_ is varied as described below.

The energy *E* from Equation 1 is sampled using a standard MC procedure (i.e., `MonteCarloMinimizer`) with temperature *T* = 10^−4^, where acceptance is defined by *a* = min [1, exp(− Δ*E*_Ω_*/T*)] with Δ*E*_Ω_ being the difference in system energy between the proposed and current configurations, i.e., `System`s from `BAGEL`. New proposals are generated through substitutions, insertions, and deletions of amino acids, with these move types chosen with probabilities 0.70, 0.15, and 0.15 respectively. When a substitution is selected, we uniformly sample the new amino acid identity from the remaining 18 possibilities (excluding the current residue and cysteines, which are excluded by default in `BAGEL`).

#### 4.2.1 Preliminary Weight Parameter Sweep

To determine an appropriate value for *w*_chemical_, we performed a preliminary sweep over five weight values: *w*_chemical_ ∈ {0.0001, 0.0005, 0.001, 0.005, 0.01}. For each enzyme and weight combination, we ran five independent MC trajectories of 100,000 steps each. From each trajectory, we extracted the last 100 unique sequences by iterating backwards from the final step, ensuring representative sampling from the equilibrated portion of the energy landscape. These sequences were folded with ESMFold, and we computed RMSD (heavy atoms) and pLDDT over the immutable region to assess structural preservation of the active site. Based on the trade-off between miniaturization and structural fidelity (Figure 2), we selected *w*_chemical_ = 0.001 for subsequent production runs.

#### 4.2.2 Production Runs

We performed 50 independent production runs of 100,000 MC steps each, carried out for each of the four enzymes. From each trajectory, we extracted unique sequences after discarding the first 25,000 transient steps, yielding a pool of candidate mini-variants for subsequent filtration (Section 4.4). The transient region refers to the energy evolution shown in Figure 1C.

### 4.3 Manual Trimming Baseline

To assess the added value of PLM-guided optimization over naive sequential deletion, we define a deterministic manual trimming baseline (Algorithm 1 in Appendix B).

### 4.4 Filtration Protocol

From the pool of unique sequences obtained from the production runs (Section 4.2.2), we fold all candidates with three structure prediction models: ESMFold [12], Chai-1 (without MSA, using ESM-2 embeddings) [14], and Boltz-2 (with MSA) [15]. We filter out all entries that have an RMSD of the immutable region above a specific cutoff value (*r*_RMSD_) for any of the folded structures. RMSD is computed using `biotite` [38].

To bias towards soluble sequences, we employ SolubleMPNN [16] and compute the log-likelihood of a sequence, using the `single_aa` approximation where the log-likelihood is computed independently for each position *i* as:

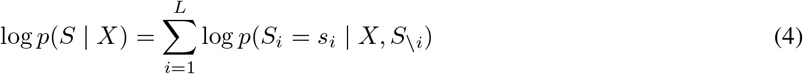

where *S* is the sequence, *X* is the protein backbone, *L* is the sequence length, *s*_*i*_ is the amino acid at position *i*, and *S* _*i*_ denotes the sequence at all positions excluding *i*.

We rank by the mean SolubleMPNN over the three folded structures, resulting in top-16 mini-variants that we consider for further validation with molecular dynamics. Note, we do not carry out any diversity filtering through sequence clustering, as we found that the final selection of the 16 mini-variants would not change dramatically, and we realize that a few point mutations could actually have a significant impact on function. All probability density distributions and kernel density contours shown in the figures were estimated using `scipy.stats.gaussian_kde` with Scott’s rule for bandwidth selection, with RMSD densities computed in log_10_-space. Table 3 summarizes the key filtration metrics.

**Table 3:** Filtration pipeline applied to miniaturized enzyme candidates. Filtration metrics for each enzyme showing the number of unique sequences after initial deduplication (25,000 transient steps), the RMSD threshold used for filtering, and the number of entries remaining after filtering entries where all three folding models (ESMFold [12], Chai-1 [14], Boltz-2 [15]) have immutable region RMSD below the threshold.

| Enzyme | # after MC | $r_{\text{RMSD}}$ (Å) | # after RMSD |
| --- | --- | --- | --- |
| petase | 405 | 1.0 | 267 |
| protease | 338 | 1.0 | 142 |
| taq | 2,847 | 1.5 | 91 |
| vioA | 1,648 | 5.5 | 19 |

### 4.5 Molecular Dynamics

We perform 500 ns of all-atom molecular dynamics simulations in explicit solvent for the top-16 mini-variant candidates per enzyme using OpenMM [39]. Prior to simulation, we process each folded structure with `pdbfixer` to correct structural issues, such as missing hydrogens. We then add hydrogens at pH 7.0 and solvate each system in a water box with 2.0 nm padding. Simulations employ the AMBER14-all force field [40] with TIP3P-FB water [41], particle mesh Ewald (PME) electrostatics, and a 2.0 nm cutoff for non-bonded interactions. Each system is minimized in three stages: (i) minimization of hydrogens with all non-hydrogen atoms restrained (1 × 10^10^ kJ/mol/nm^2^), (ii) minimization of water with the protein restrained (1 × 10^10^ kJ/mol/nm^2^), and (iii) minimization of the full system without restraints. We then equilibrate at 300 K for 10 ns, followed by a 500 ns production run with a 2 fs timestep. Trajectories are saved every 100 ps, recording only protein atoms. Trajectory analysis was done using `MDAnalysis` [42, 43]. We note that all simulations were performed on the *apo* protein without cofactors; in particular, the FAD cofactor required by VioA was not included, and thus the MD results for *vioA* should be interpreted with this caveat in mind.

## 5 Code and Data Availability

The code is available in the `scripts/mini-enzymes` folder in the GitHub repository of `BAGEL` [10] package (https://github.com/softnanolab/bagel). All of the data has been deposited to Zenodo (https://doi.org/10.5281/zenodo.18854113).

## 6 Acknowledgements

We thank Erik Baldauf, Toby Hallett, and Akashaditya Das for insightful discussions. J.L. acknowledges the President’s PhD scholarship at Imperial College London for funding. We acknowledge computational resources and support provided by the Imperial College Research Computing Service (http://doi.org/10.14469/hpc/2232). We are grateful to the UK Materials and Molecular Modelling Hub for computational resources, which is partially funded by EPSRC (EP/T022213/1, EP/W032260/1 and EP/P020194/1). This work made use of the Isambard-AI service at the University of Bristol. Isambard-AI is funded by the UK Government through the Department for Science, Innovation and Technology (DSIT) and delivered in partnership with the University of Bristol and HPE/Cray. This research was also supported by grants from NVIDIA and utilized NVIDIA’s GPU access through the Academic Grant for Simulation and Modeling. We thank Modal for computational support through their academic grants program.

## 7 Author Contributions

S.A-U. designed the research. J.L. implemented the research. H.A. secured funding. F.D. performed preliminary research and analysis. J.L., H.A., J.W. and S.A-U. analyzed the data and wrote the manuscript.

## A Chemical Potential Energy Weights Sweep

**Figure 7:**
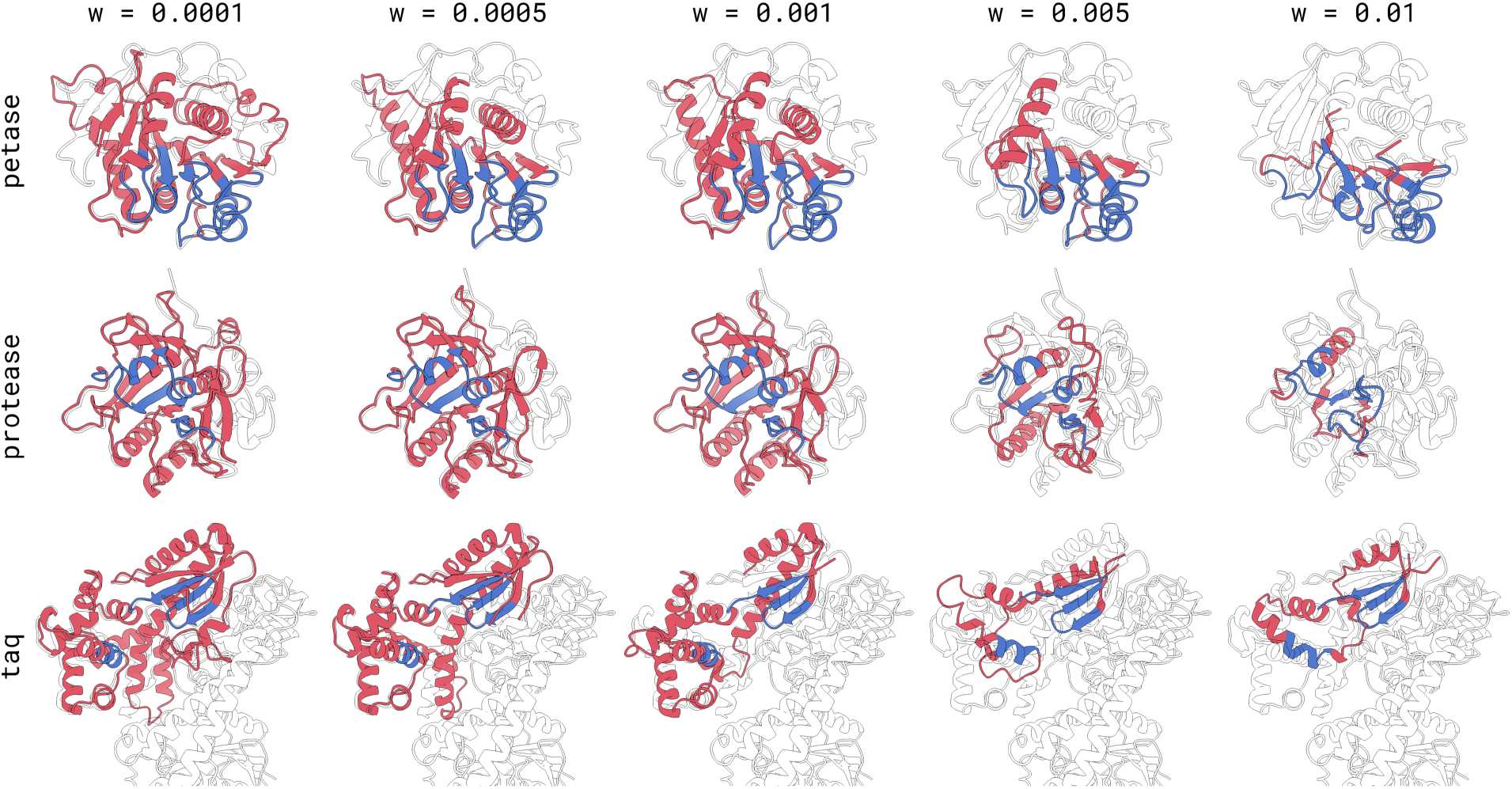
Structural impact of chemical potential weights on enzyme miniaturization. Structural representations of mini-variant sequences for *petase* (first row), *protease* (second row), and *taq* (third row) across different weights *w*_chemical_ of the sequence length penalty. The immutable region is highlighted in blue, the full-length WT sequence is shown in the black outline. All structures are folded with ESMFold [12] (including the WT). As *w*_chemical_ increases from left to right, the sequences become progressively shorter, and the immutable region becomes structurally compromised.

## B Manual Trimming Algorithm

### Algorithm 1

Manual Trimming Baseline

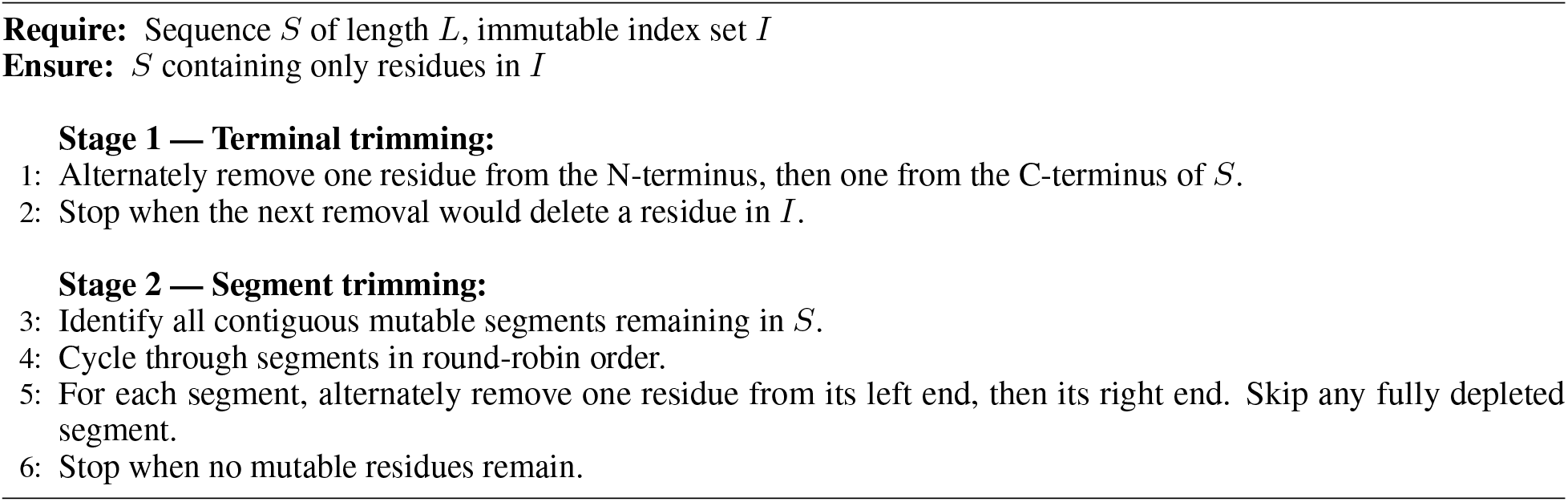

## C RMSD vs. pLDDT Analysis

**Figure 8:**
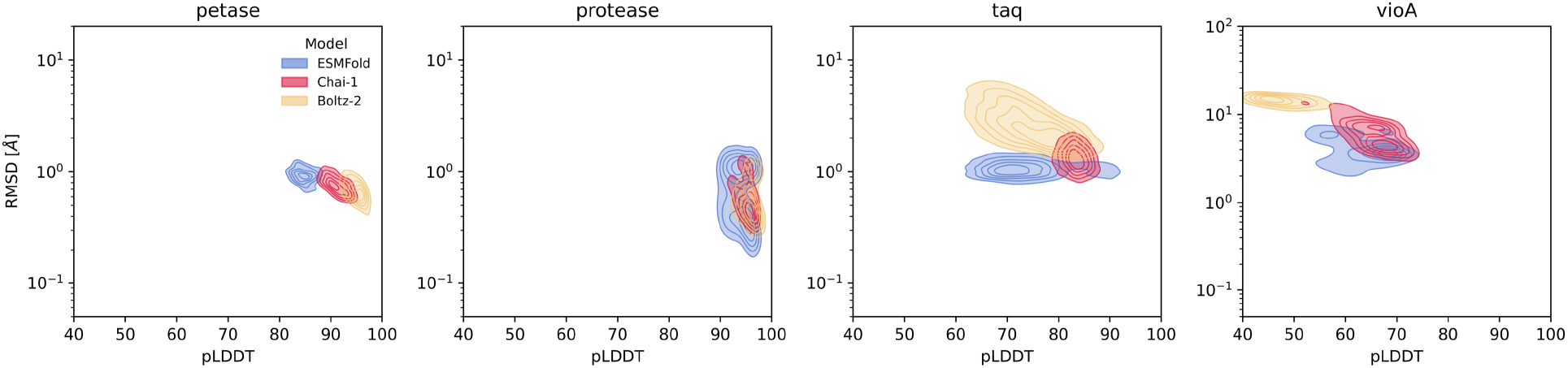
Joint RMSD–pLDDT analysis of miniaturized enzyme variants. Comparison of RMSD and pLDDT densities for mini-variant sequences across four enzymes. Kernel density estimation contours are shown for structures folded using ESMFold [12] (blue), Chai-1 [14] (red), and Boltz-2 [15] (yellow). Densities were estimated using `scipy.stats.gaussian_kde` with Scott’s rule for bandwidth selection in (pLDDT, log_10_(RMSD)) space. Contour levels represent normalized density from 0.2 to 1.0 of the per-model maximum. In general, lower RMSD in the immutable region to the WT corresponds to higher pLDDT.

## D Solubility and Hydrophobicity Analysis

To assess the solubility and hydrophobicity of the mini-variants before expressing them in a lab, we employed the SolubleMPNN model [16] to rank sequences by predicted solubility. On top of that, we also computed the largest hydrophobic patch area (adapted from BoltzGen [44]). Figure 9 summarizes the distributions of these two metrics, always comparing them to the WT.

For the largest hydrophobic patch area, we first selected all carbon atoms belonging to hydrophobic residues (Val, Ile, Leu, Phe, Met, Trp) and computed the per-atom solvent-accessible surface area (SASA) using the Shrake-Rupley algorithm [45] as implemented in `biotite` [38], with a water probe radius of 1.4 Å. Only surface-exposed atoms (SASA *>* 0) were retained. These surface hydrophobic carbons were then clustered spatially using `DBSCAN` [46] with *ε* = 5.0 Å and `min_samples` = 1, grouping proximal atoms into contiguous patches. The largest hydrophobic patch area is defined as the sum of SASA values of all atoms in the largest cluster:

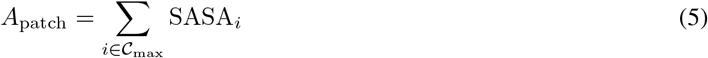

where 𝒞_max_ is the largest `DBSCAN` cluster.

In terms of the largest hydrophobic patch area, we instead observe an opposite trend, and most of our mini-variants have smaller patches than the WT. We note, however, that we do not normalize the patch area to the overall size of the protein, which might explain the difference. Whether the relative patch size to the overall size of the protein, or the absolute patch size, is the more relevant metric is an open research question, and further experimental validation might elucidate the answer.

Overall, this yields only heuristics, and thus we only rely on the SolubleMPNN log-likelihood as a minor biasing technique to get higher likelihood sequences; rather than using it as a strict filtering criterion, or for that matter, any of the other metrics. All in all, these show that despite not explicitly optimizing for solubility, the PLM-based approach is able to retain this property in some cases, just by constraining the embeddings of the active site residues.

Despite this, under the assumption that SolubleMPNN log-likelihood is a good proxy for solubility, we identify two routes for improving predicted solubility in future work: (a) integrating SolubleMPNN as a `MutationProtocol` into `BAGEL` to guide optimization toward more expressible sequences, or (b) applying a post-hoc “repainting” step where the mutable regions are redesigned using inverse folding models conditioned on the miniaturized backbone, following standard structure-based protein design workflows.

**Figure 9:**
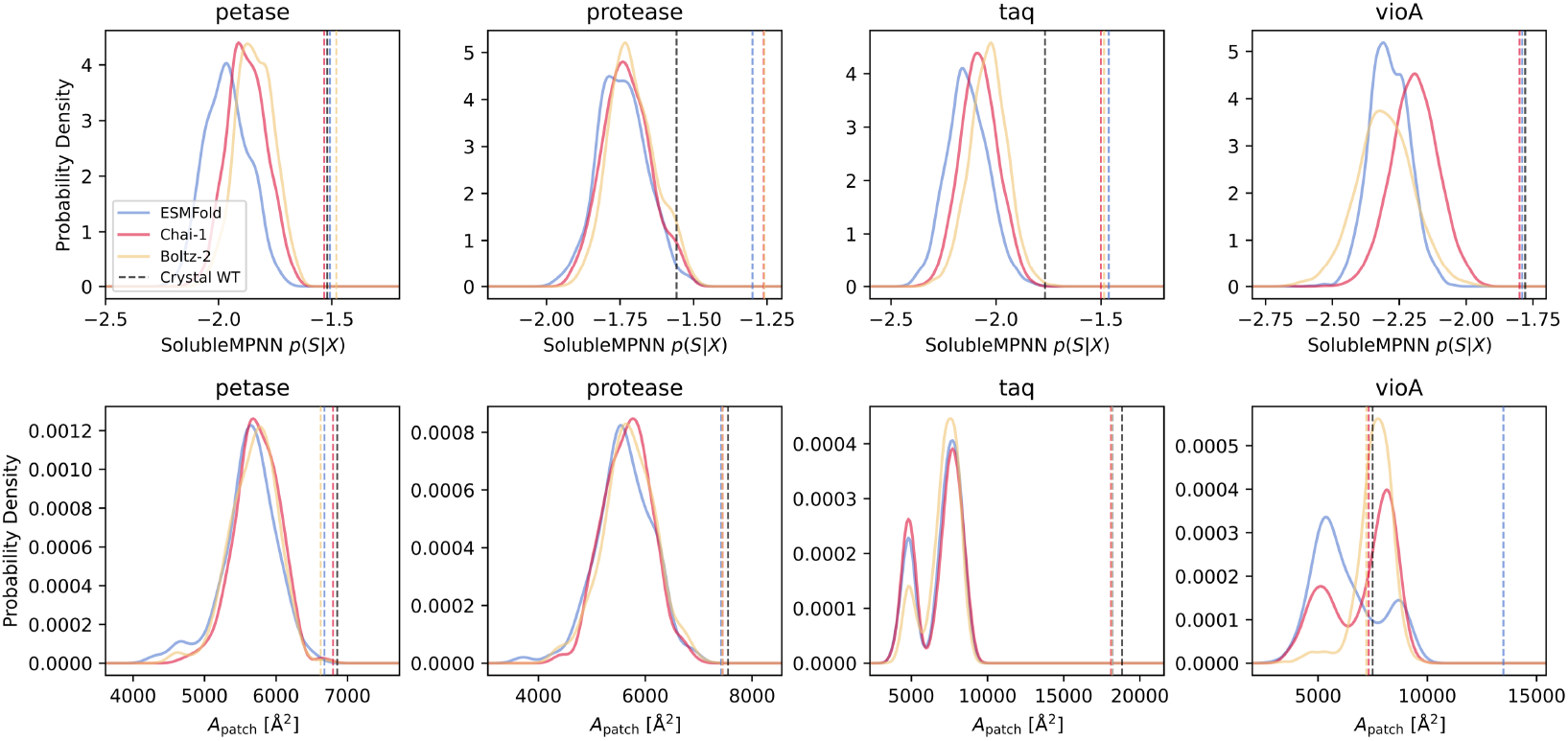
Computational solubility assessment of miniaturized enzyme variants. Distributions of two solubility and hydrophobicity metrics for mini-variant sequences across four enzymes. Top row: SolubleMPNN [16] log-likelihood values *p*(*S* | *X*) (defined in Equation 4). Bottom row: Largest hydrophobic patch area computed via Shrake-Rupley solvent-accessible surface analysis (adapted from BoltzGen [44] and defined in Equation 5). Probability density distributions are shown for sequences folded with ESMFold [12] (blue), Chai-1 [14] (red), and Boltz-2 [15] (yellow). Vertical dashed lines indicate wild-type scores for each folding model and the crystal structure (black). Mini-variant sequences show lower SolubleMPNN log-likelihood values than wild type, likely reflecting their novelty rather than definitive insolubility.

## E Molecular Dynamics Data

**Figure 10:**
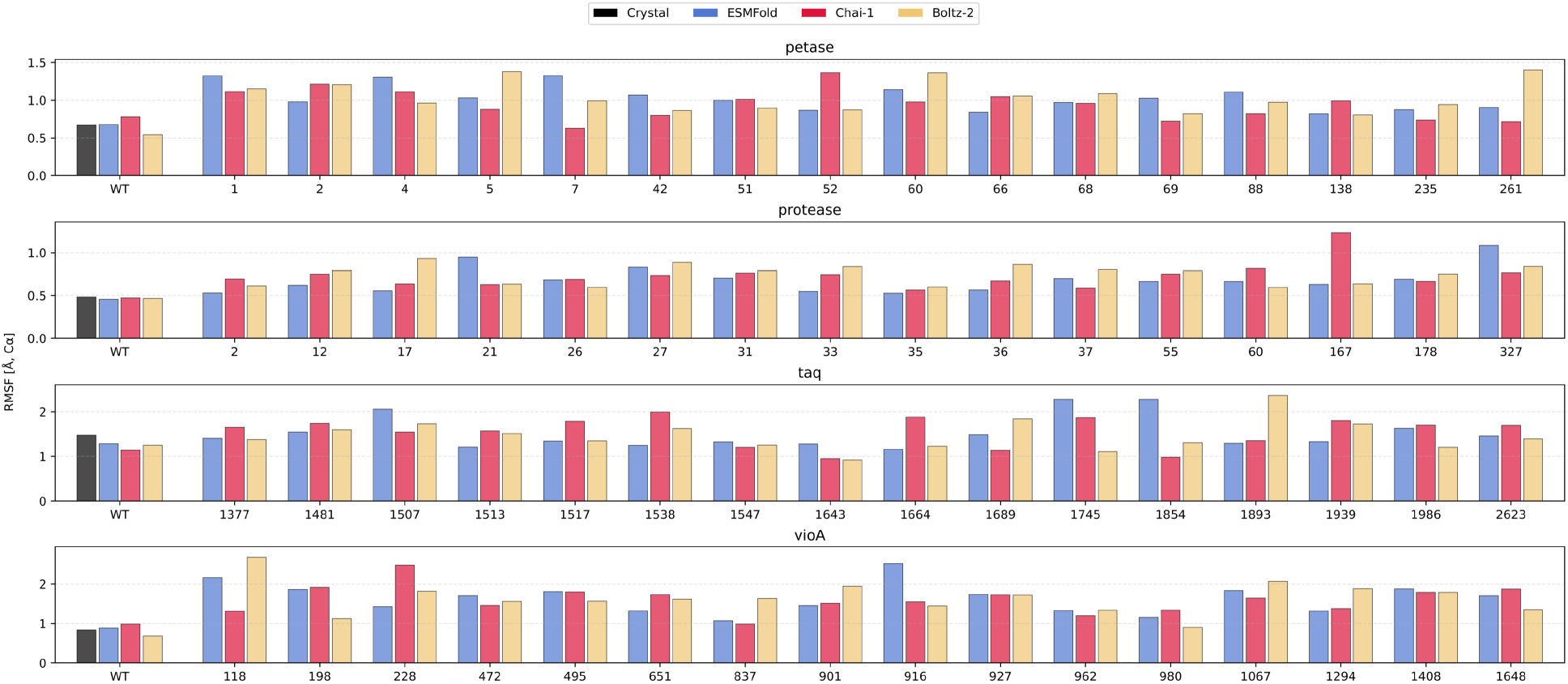
MD-derived RMSF for all miniaturized enzyme candidates. RMSF (defined in Equation 2) for residues *i* ∈ *I*, where *I* is the immutable residue group (defined in Equation 1), computed from molecular dynamics simulations for mini-variant sequences across four enzymes. RMSF values are shown for the wild-type (WT) and selected mini-variants, with starting configurations folded using ESMFold [12] (blue), Chai-1 [14] (red), and Boltz-2 [15] (yellow). The WT crystal structure reference is shown in black. Mini-variants show variable RMSF values across different designs, as well as among different folding models. In general, WT shows lower values for *vioA* and *petase*, with similar fluctuations or sometimes even higher for *taq* and *protease*.

**Figure 11:**
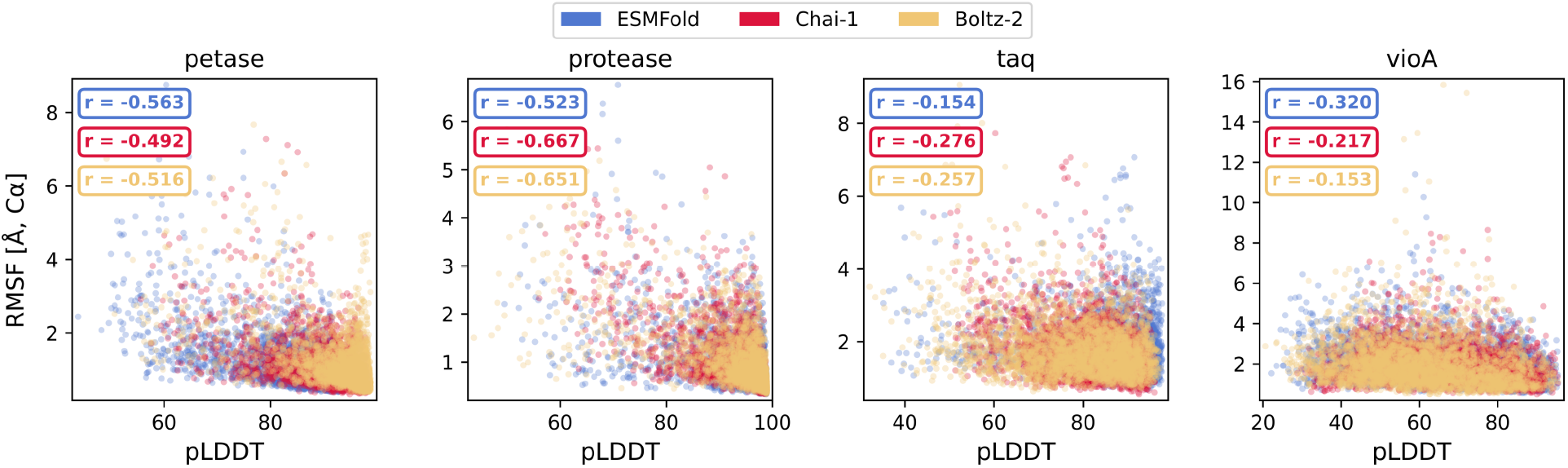
Correlation between structure prediction confidence and thermal fluctuations for miniaturized enzyme variants. Correlation between structure prediction confidence (pLDDT) and thermal fluctuations (RMSF, defined in Equation 2) for the four enzymes. Points are colored by the folding model used: ESMFold [12] (blue), Chai-1 [14] (red), and Boltz-2 [15] (yellow). Pearson correlation coefficients (r) are displayed for each enzyme-model combination. The correlation strength varies by enzyme, with *petase* and *protease* showing the strongest relationships, while *taq* and *vioA* exhibit weaker correlation. This provides supporting evidence that the pLDDT score can be used as a proxy for thermal fluctuations in further optimization steps of the design campaign.

